# The fitness burden imposed by synthesising quorum sensing signals

**DOI:** 10.1101/050229

**Authors:** A. Ruparell, JF. Dubern, CA. Ortori, F. Harrison, NM. Halliday, A. Emtage, M. Ashawesh, CA. Laughton, SP. Diggle, P. Williams, DA. Barrett, KR. Hardie

## Abstract

It is now well established that bacterial populations utilize cell-to-cell signaling (quorum-sensing, QS) to control the production of public goods and other co-operative behaviours. Evolutionary theory predicts that both the cost of signal production and the response to signals should incur fitness costs for producing cells. Although costs imposed by the downstream consequences of QS have been shown, it has not been demonstrated that the production of QS signal molecules (QSSMs) results in a decrease in fitness. We measured the fitness cost to cells of synthesising QSSMs by quantifying metabolite levels in the presence of QSSM synthases. We found that: (i) bacteria making QSSMs have a growth defect that exerts an evolutionary cost, (ii) production of QSSMs correlates with reduced intracellular concentrations of QSSM precursors, (iii) the production of heterologous QSSMs negatively impacts the production of a native QSSM that shares common substrates, and (iv) supplementation with exogenously added metabolites partially rescued growth defects imposed by QSSM synthesis. These data provide the first direct experimental evidence that the production of QS signals carries fitness costs to producer cells.

**Originality-Significance Statement:** Bacterial cells within populations communicate with each other to control social behaviors by producing diffusible quorum sensing (QS) signal molecules. Evolutionary theory predicts that both the cost of signal production and the response to signals should incur fitness costs for producing cells. Here we provide the first empirical evidence that the production of QS signals incurs fitness costs to producing cells. Since QS plays a major role in bacterial pathogenicity, this finding will underpin novel antimicrobial strategies that are urgently needed to replace currently available antimicrobials that are becoming obsolete through the ever-rising incidence of resistance.

## Introduction

Communication systems are widespread in plants, animals and microorganisms. For true communication (signaling) to evolve, signals must transfer information that benefits both the signaller and the receiver. Whether signals are visible, acoustic or chemical in nature, their production implies a cost to the emitter, but these costs are often difficult to measure experimentally (Smith & Harper, 2003; Keller & Surette, 2006; Diggle *et al.*, 2007b; Popat *et al.*, 2015). Many bacterial species communicate using small diffusible signals to co-ordinate social behaviours in a process termed quorum sensing (QS) (Atkinson & Williams, 2009; Darch *et al.*, 2012). QS signaling molecules (QSSMs) are synthesized inside the bacterial cell and released into the surrounding environment. Once accumulated to a threshold concentration, the QSSMs drive the expression of genes encoding public goods and other social behaviors that benefit the surrounding population of cells. Previous work has shown that there are fitness costs associated with producing QS regulated public goods, and that these costs are significant enough for non-producing cheats to evolve and spread in populations (mutants that can respond to QSSMs, but do not make them) (Diggle *et al.*, 2007a; Rumbaugh *et al.*, 2009; Pollitt *et al.*, 2014). Evolutionary theory has also predicted that the production of the QSSMs themselves should also incur a fitness cost, but there have not been any studies explicitly measuring the fitness consequences of producing them. Any fitness costs are presumed to be a drain in metabolites, and Keller and Surrette estimated that production of each of three well-studied QSSM classes (oligopeptides, *N*-acylhomoserine lactones (AHLs) and Autoinducer-2 (AI-2) impose a metabolic cost of 184 ATPs, 8 ATPs and 0-1 ATP respectively (Keller & Surette, 2006). We therefore set out to experimentally determine whether there are metabolic consequences for QSSM synthesis.

We chose to work with the two dedicated QSSM LuxI-type synthases (Lasl and Rhll) that produce AHLs in the opportunistic, multi-antbiotic-resistant pathogen *Pseudomonas aeruginosa*. LasI, synthesizes long chain AHLs (predominantly *N*-(3-oxododecanoyl)-L-homoserine lactone (OC_12_-HSL)). Rhll primarily synthesizes the short chain AHL (*N*-butanoyl-L-homoserine lactone (C_4_-HSL)). *P. aeruginosa* incorporates these AHL synthases into a complex hierarchical QS network which controls virulence factor production and thus pathogenicity in plants and animals (including humans) (Williams & Cámara, 2009). Lasl and and Rhll were each introduced into a heterologous host, *Escherichia coli* that does not naturally produce AHLs. We reasoned that it is important to determine whether there are fitness costs specifically associated with QSSM production *per se*. A future challenge remains with respect to understanding the fitness burden of a complete QS system in its natural, adapted host. In this context, the added cost of responding to the QSSM which may in turn be integrated into a complex regulatory network or have pleiotropic effects unlinked to the response to signals can also be considered.

Synthesis of AHLs depends on the availability of the precursors: an appropriately charged acyl carrier protein (acyl-ACP) and *S*-adenosyl-L-methionine (SAM) (More *et al.*, 1996; Jiang *et al.*, 1998; Parsek *et al.*, 1999; Raychaudhuri *et al.*, 2005). The donation of the acyl group from acyl-ACP to the amine of SAM results in the formation of the AHL and release of 5’-methylthioadenosine (MTA) (More *et al.*, 1996; Jiang *et al.*, 1998; Parsek *et al.*, 1999). Both Lasl and Rhll use SAM, but link it to a different fatty acid. We can assume they are approximately catalytically equivalent, although kinetic data is only available for Rhll (Jiang *et al.*, 1998; Raychaudhuri *et al.*, 2005). As the major methyl donor in eubacterial, archaebacterial and eukaryotic cells, SAM is a critically important metabolite (Cooper *et al.*, 1993; Low *et al.*, 2001). The availability of SAM and relative flux through the AMC (activated methyl cycle: to which SAM contributes see supplementary Figure 1) significantly impacts upon central metabolism and influences cell fitness, as has been well documented in the context of two AMC enzymes, LuxS and Pfs (Winzer *et al.*, 2003; Vendeville *et al.*, 2005; Hardie & Heurlier, 2008; Heurlier *et al.*, 2009; Doherty *et al.*, 2010; Halliday *et al.*, 2010).

Here we test the metabolic (by measuring AMC-linked metabolite levels) and fitness (by monitoring growth) costs of making an AHL QSSM in the heterologous host, *E. coli*. We show that (i) signal production causes a growth defect that imposes a fitness disadvantage in mixed populations, (ii) signal production correlates with reduced intracellular concentrations of the substrates required, (iii) heterologous signal production negatively impacts native signal production, (iv) supplementation with exogenously added metabolites partially rescued growth defects imposed by signal synthesis. Our findings demonstrate that the fitness cost of generating the QS signals required for co-ordinated social behaviour in bacteria can be substantial.

## Experimental Procedures

### Bacterial strains and growth conditions

Strains and plasmids used in this study (supplementary Table 1) were routinely grown in Luria-Bertani medium (LB) or on nutrient agar plates at 37°C. A MOPS minimal medium (MMM) was prepared as described previously (Vendeville *et al.*, 2005). Antibiotics were added at the following concentrations: carbenicillin 25 μg/ml; tetracycline 25 μg/ml and 100 μg/ml kanamycin. Isopropylthio-β-D-galactoside (IPTG) was added at a final concentration of 1 mM, unless otherwise indicated. Growth was followed by estimating optical densities at 600 nm using a 1 in 10 dilution of cultures into the growth medium to ensure accurate spectrophotometer readings, or using viable cell counts (colony forming units: CFU).

### DNA manipulation and cloning procedures

DNA was purified using a plasmid purification kit (Qiagen) or Wizard genomic DNA purification kit (Promega). Restriction enzyme digestion, ligation and agarose gel electrophoresis were performed using standard methods (Sambrook *et al.*, 1989). Restriction fragments were routinely purified from agarose gels using a QIAquick kit (Qiagen). Transformation of *E. coli* was carried out by electroporation (Farinha & Kropinski, 1990). Oligonucleotide primers (supplementary Table 1) were synthesised by Sigma Genosys. Both strands of cloned PCR products were sequenced by the DNA Sequencing Laboratory at the University of Nottingham (United Kingdom). Nucleotide and deduced amino acid sequences were aligned using Clustal W (http://clustalw.genome.jp/).

### Plasmid construction and site directed mutagenesis

The 606-bp *lasl* and *rhll* genes were PCR-amplified using chromosomal DNA of *P. aeruginosa* PA01 as the template and the primer pairs *lasl* Forward/*lasl* Reverse or *rhll* Forward/*rhll* Reverse respectively. The purified 0.624 kb fragments were ligated into the pGEM-T Easy vector (designated pGEMT-*lasI* or pGEMT-*rhll*) and released with *Eco*RI-*St*uI digestion for cloning into shuttle vector pME6032 (pME-*lasI* and pME-*Rhll)*.

To mutate the *lasl* and *rhll* genes, degenerated phosphorylated PCR primer pairs amplified pGEMT-*lasI* or pGEMT-*rhll*. Following recircularisation and transformation into DH5α, the *lasl* and *rhll* mutant fragments were recovered from pGEMT-Easy and inserted as *Eco*RI-*St*uI fragments into the pME6032 vector. The point mutations were confirmed by sequencing.

### SDS-PAGE and western blotting

This was undertaken as described previously (Cooksley *et al.*, 2003). Anti-Rhll was diluted to 1:2000 and anti-mouse IgG HRP (Sigma) to 1:1000.

### Small molecule analysis

AHLs were quantified by LC-MS/MS as described previously (Ortori *et al.*, 2011). For extraction of intracellular AMC metabolites, bacteria were grown in 125 ml IPTG supplemented LB in 500 ml Erlenmeyer flasks at 37^°^C for 12 h. Samples of 5 ml containing equivalent cell densities for each strain were quenched with 15 ml of phosphate-buffered saline (PBS), cells were lysed, metabolites derivatized and detected (Halliday *et al.*, 2010). To determine intracellular MTA levels, the existing protocol (Halliday *et al.*, 2010) was modified to use 100% MeOH for extraction and samples were reconstituted in 100 μl of dH_2_0 and analysed by liquid chromatography-tandem mass spectrometry using a Quattro Ultima (Waters Micromass, Manchester, UK) in conjunction with an Agilent 1100 LC system (Agilent Technologies, Waldbron, Germany) with a cooled autosampler [30]. HPLC was carried out using a Synergi 4u Hydro-RP (4 μm, 150 _ 2.0 mm, Phenomenex, Macclesfield, UK) with a guard column fitted. AI-2 was quantified by (Heurlier *et al.*, 2009) and expressed as the change in bioluminescence of the reporter strains (bioluminescence in the presence of extract/background bioluminescence in the presence of sterile medium).

### Mixed Population Competition Assay

Approximately 10^5^ cells from overnight cultures of each strain or a 1:1 mixture of two strains were inoculated into 2ml MMM + tetracycline. Four wells of each population were supplemented with IPTG and four left uninduced. Cultures were grown shaking for 24 h at 37°C. An aliquot of each population was diluted and plated onto two LB + tetracycline agar plates for colony counting. To calculate the final proportion of each strain in the mixed populations, 50 colonies from each mixed population were randomly selected and streaked across an aliquot of the *E coli* biosensor pSB1075 (Winson *et al.*, 1998) on LB + tetracycline + IPTG agar plates. The biosensor carries a *lux* reporter that is expressed in the presence of OC_12_-HSL, thus cross-streaking with *E. coli* MG1655(pME-*lasI*) produces luminescent streaks after overnight growth.

A Hamamatsu Aequoria darkbox and M4314 Image Intensifier Controller were used along with the software Wasabi 1.5 to image plates and score numbers of light and dark streaks. To calculate the false negative and false positive rates, ten colonies from each pure population were also streaked across the biosensor and imaged after overnight growth. Bayes’ theorem was applied to these data to calculate the probability that a light streak was *E. coli* MG1655(pME-*lasI*) and that a dark streak was *E. coli* MG1655(pME6032); these were 0.99 and 0.94 respectively and we adjusted observed light/dark numbers in the mixed populations to take account of this.

The evolutionary fitness of *E. coli* MG1655(pME-*lasI*) relative to *E. coli* MG1655(pME6032) in mixed populations was calculated using:

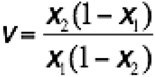

where *X*_1_ and *X*_2_ are the initial and final frequencies of *E. coli* MG1655(pME-*lasI*) in the population, respectively (Winson *et al.*, 1998). When the two genotypes have equal fitness, *X*_1_ = *X*_2_ and *v* = 1. Values of *v* < 1 reflect being outcompeted by MG1655(pME6032) and values > 1 indicate that MG1655(pME-*lasI*) outcompetes MG1655(pME6032). The relative fitness of *E. coli* MG1655(pME-*lasI*) in pure culture was calculated by randomly pairing pure MG1655(pME-*lasI*) and MG1655(pME6032) populations with IPTG treatment and applying the same formula. Data were analysed using ANOVA in R 2.14.0 (R Development, 2011). Total population size and relative fitness were both square root transformed to meet the assumptions of parametric tests and when dropping of an outlier in the growth data caused loss of orthogonality the *car* package (Fox & Weisberg, 2011) was used to implement ANOVA with Type II sums of squares.

## Results

### Production of chemical signals used for social communication compromises cell fitness

Bacterial populations can communicate using QSSMs, and it has been shown that QS responses impose a fitness cost (Pai *et al.*, 2012). To test whether QSSM synthesis is also metabolically costly, the genes encoding QSSM synthases Lasl and Rhll were expressed from the shuttle vector pME6032 in *E. coli*. Having hypothesized that QSSM synthesis would compromise cell fitness, the growth profiles of the strains were compared in both minimal (MMM; Figure 1) and rich (LB; supplementary Figure 4e and Figure 5e) media. The empty plasmid pME6032 had no detrimental effect on growth (referred to as non-producer). In LB medium containing abundant nutrients, there was a slightly slower initial growth rate and lower final density upon induction of the AHL synthase LasI demonstrating a fitness cost and by extension a potential metabolic cost (supplementary Figure 4e). In line with observations that disruption of one of the metabolic cycles feeding into AHL synthesis (the AMC) is more readily reflected in growth defects in defined media limited for the sulphur sources that feed into this pathway (Winzer *et al.*, 2003; Doherty *et al.*, 2006; Heurlier *et al.*, 2009; Holmes *et al.*, 2009; Doherty *et al.*, 2010), more drastic effects on growth were observed in MMM. The RhlI-producer MG1655(pME-*rhlI*) did not grow as well as the empty vector control initially (Figure 1), but over time achieved the same final population density. In accordance with the higher level of LasI production (supplementary Figure 2a,b), it had an even more dramatic effect on growth, with the LasI-producer MG1655(pME-*lasI*) growing much slower than the non-producer, and failing to achieve a population size (OD_600_) equivalent to the other strains within 22 h in MMM (Figure 1).

**Fig. 1.**
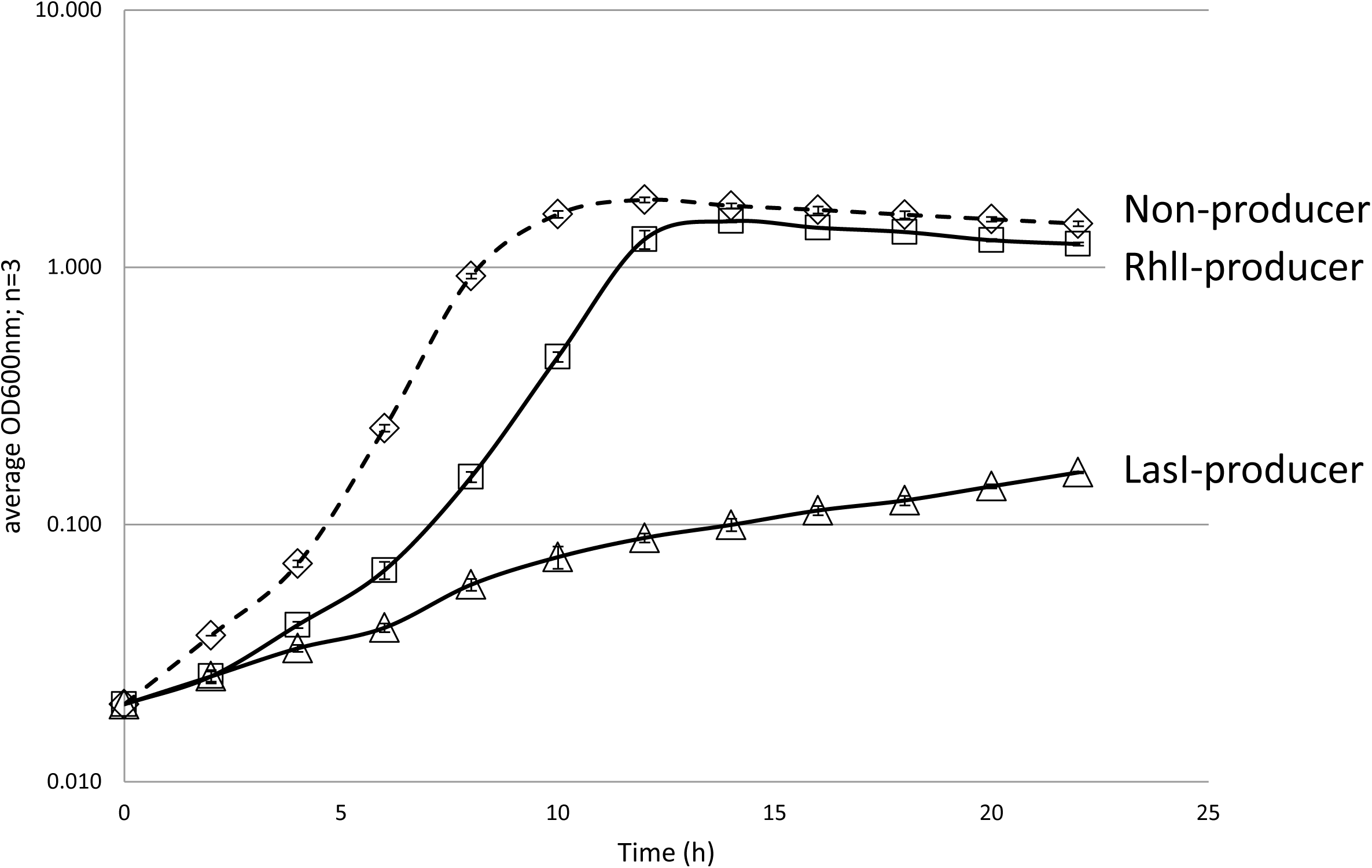
QSSM synthesis alters the fitness of bacteria. Three *E. coli* strains non-producer MG1655(pME6032), LasI-producer MG1655(pME-*lasl*), and Rhll-producer MG1655(pME-*rhlI*) were inoculated into 125 ml MMM + tetracycline + IPTG and grown with shaking at 37 °C in 500 ml.Optical density at 600 nm and the data are indicated as means ± standard deviations for three independent cultures.

The production of the major cognate QSSMs in *E. coli* culture supernatants by Lasl and Rhll (OC_12_-HSL and C_4_-HSL respectively) was confirmed by thin-layer chromatography (TLC) (data not shown) with quantification using sensitive LC-MS/MS (supplementary Figure 2c,d). In line with an absence of inhibition of *E. coli* growth inflicted by exogenous addition of these concentrations of QSSMs (supplementary Figure 6; up to 800 μM), no defect in growth was observed in rich medium with the Rhll-producing strain (supplementary Figure 5e), and the growth of the LasI-producer was only marginally reduced in rich media (supplementary Figure 4e). The masking of the overall cost of producing signals by growth in rich media suggests that it could at least in part derive from an energetic cost as it can be topped up by provision of exogenous metabolites. To determine the lower limit for the level of QSSM production that results in a cell fitness cost, IPTG concentrations were titrated down to reduce the amount of Rhll or Lasl produced. At the point where QSSMs were barely detectable, all strains grew at a similar rate to reach a comparable stationary phase population density.

### Mutations that prevent signal production rescue bacterial growth defects

It is possible that the observed growth defects may be due to the burden of protein overproduction in producers, rather than signal synthesis. To discount this, several mutants were constructed. Whilst some Lasl/Rhll mutants maintained the ability to synthesize AHLs, others did not, and only those able to make AHLs reduced the growth of *E. coli* (supplementary Figure 4 and 5).

The structures of the AHL synthases LasI and EsaI were used to model the predicted structure of Rhll (supplementary Figure 3a) in order to identify key residues predicted to be involved in catalysis as targets for parallel site directed mutagenesis of Lasl and Rhll. The catalytic residues chosen for mutagenesis were based on those identified in a previous study (Parsek *et al.*, 1997) that screened the activity of a collection of mutations in Rhll. Residues F27 and W33 of LasI are important for SAM binding, and S103 appears to participate within a cluster of other residues to maintain tertiary structure interactions and may also perform a catalytic function (Hoang & Schweizer, 1999). Changes were also made to residues that may alter a specific property of the active site (see supplementary Table 2), and to R23 of LasI since we predicted it would be involved in catalysis. All the residues selected mapped to the vicinity of a pocket hypothesised to be the active site (supplementary Figure 3b).

The AHL synthase mutants that could be overproduced as a protein of the predicted size were selected for further study (supplementary Figure 4c, 5c). Wild type levels of QSSMs were produced by the LasI mutants F27Y and S103A (supplementary Figure 4a). Similarly, mutation of S103 to either A or V in Rhll did not significantly lower total AHL production (supplementary Figure 5a). With the exception of LasI F27L, the other producer mutants completely lost the ability to synthesise AHLs. All the mutants able to synthesize AHLs, except LasI S103A, did so in relative proportions that resembled the profile of the wild type AHL synthase. Significant concentrations of C_4_-HSL, which were not observed with wild type LasI, accounted for 20% of the AHLs made by LasI S103A (supplementary Figure 4d and 5d).

Importantly, only producer mutants that retained the ability to synthesize AHLs inhibited growth, indicating that the enzymatic activity of the QSSM synthases resulted in a fitness burden (supplementary Figure 4e,f, 5e/f). In addition, exogenous QSSMs up to 800 μM did not affect growth of the host. *E. coli* (supplementary Figure 6a,b).

### Increased levels of signal correlate with reduced intracellular concentrations of substrates used to make them

As producers synthesize AHLs from specific metabolic substrates, we hypothesized that the growth defect observed may arise as a consequence of introducing metabolite-consuming enzymes. One of the metabolites central to the AMC pathway, SAM, is a substrate for AHL synthesis. We therefore determined the profiles of the AMC metabolites (SAM, SAH, SRH, HCY and MET) to test our hypothesis. In late exponential phase cells, the level of each AMC metabolite measured was reduced following signal production (Figure 2a). Metabolite concentrations were more dramatically reduced in LasI-producers than in Rhll-producers. The metabolite which exhibited the greatest percentage concentration change (97% in Lasl-producers, 45% in Rhll-producers) was SAM (Figure 2a, supplementary Table 4). Enzyme activity was key to these metabolic perturbations because no fall in metabolite levels was observed in a mutated producer lacking the ability to synthesize AHLs (supplementary Figure 4b, 5b).

**Fig. 2.**
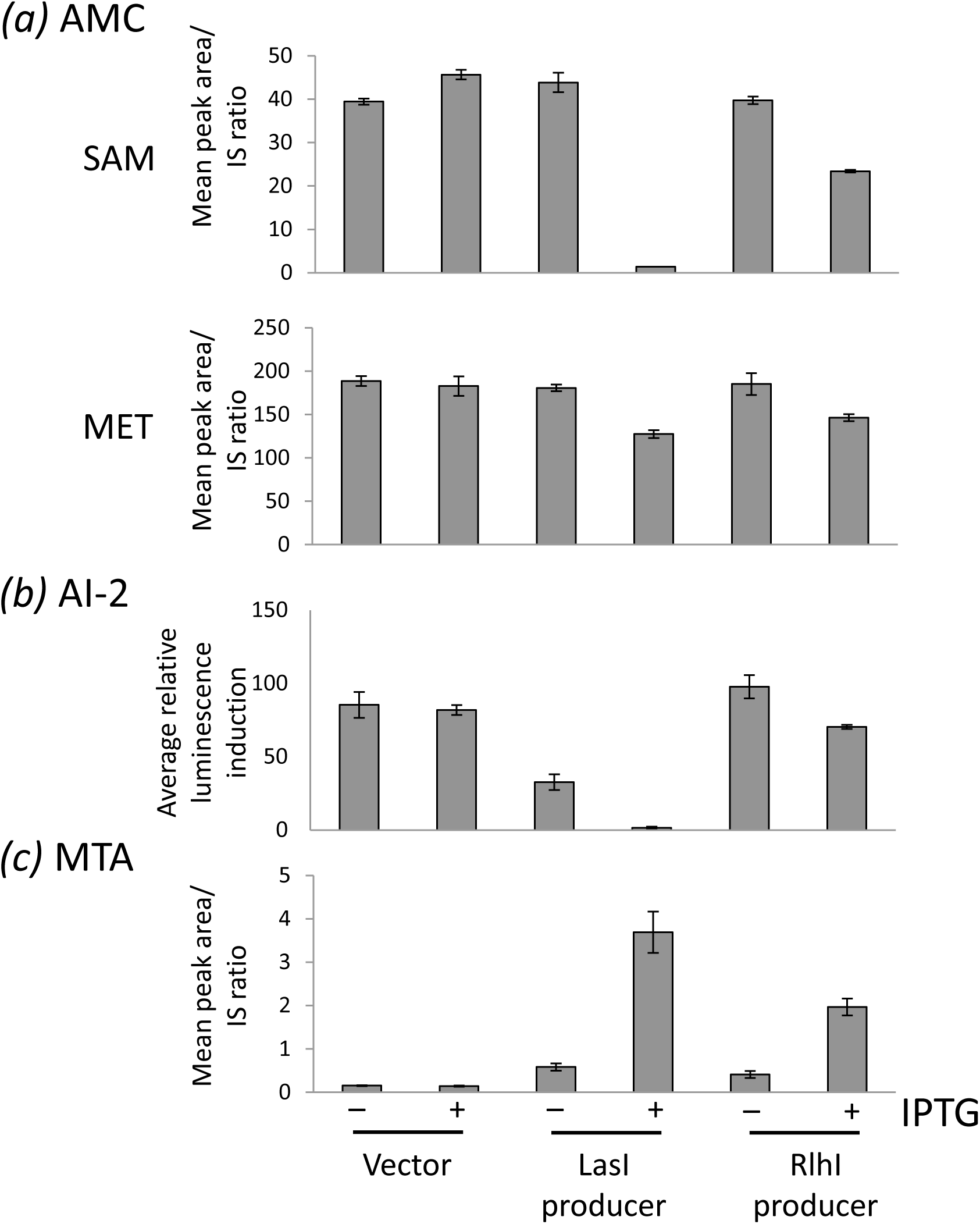
AHL synthases reduce levels of AMC-linked metabolites including the native QSSM AI-2, whilst intracellular MTA levels increase. **Panel** (*a*) Metabolite levels were determined in three *E. coli* strains MG1655(pME6032), MG1655(pME-*lasl*) and MG1655(pME-*rhlI*) grown in LB + tetracyline with and without IPTG. Intracellular accumulation of SAM, SAH, SRH, HCY and MET were determined by LC-MS analysis. SAM and MET levels represent the most and least profound modulation respectively. Peak area corresponding to each compound was divided by the peak area of the appropriate internal standard (IS) for normalisation. **Panel (*b*)** Intracellular accumulation of MTA in similarly grown MG1655Δ*pfs*(pME6032), MG1655Δ*pfs* (pME-*lasI*) and MG1655Δ*pfs* (pME-*rhl)*. **Panel c)** Spent culture supernatants were prepared from MG1655(pME6032), MG1655(pME-*lasI*) and MG1655(pME-*rhlI*) grown in LB + tetracylcine with and without IPTG until an OD_600_ of 0.75, 0.80 and 0.88. The average bioluminescence induced by AI-2 reporter *V. harveyi* strain after 2 h incubation is shown. Metabolite levels for a single experiment are shown, although the experiment has been repeated three times with similar results.

### The production of foreign signals negatively impacts the production of a native signal

One of the reactions of the *E. coli* AMC is catalysed by LuxS, and leads to the generation of AI-2. AI-2 acts as a QSSM, e.g. to stimulate the production of bioluminescence by *Vibrio harveyi*. In *E. coli*, inactivation of *luxS* has a pleiotropic effect. It is not clear what signalling role LuxS plays due to variations in strains and mutagenic strategies, although it can influence the virulence of pathogenic *E. coli* (Haigh *et al.*, 2013; Palaniyandi *et al.*, 2013). To determine the influence of a foreign QSSM synthase upon the production of a native QSSM, the amount of AI-2 produced in culture supernatants was measured. The synthase genes caused a reduction in the levels of AI-2 detected, with AI-2 levels falling by 12% in Rhll-producers and by a massive 54-fold in LasI-producers (Figure 2b).

### The fitness cost of QSSM synthesis is partly due to the production of toxic side products

AHL-synthase catalyzed production of AHLs generates a second product, MTA, which could potentially have metabolic consequences since MTA is a potent feedback inhibitor of polyamine biosynthesis (Dante *et al.*, 1983). To determine if MTA accumulates as a result of AHL synthesis, intracellular accumulation of MTA was monitored (Figure 2c). These measurements were conducted in a defined *E. coli* Δ*pfs* mutant in parallel with MG1655 because in addition to catalysing the detoxification of SAH to SRH in the reaction preceding LuxS in the AMC, Pfs can act as an MTA nucleosidase (Cornell & Riscoe, 1998), and thus potentially degrade MTA faster than we can measure it. Although MTA may not accumulate to detectable levels in MG1655, if the Δ*pfs* mutant were to show higher levels there would be the potential for transient MTA accumulation that could impact upon cell fitness. As predicted, MTA levels in *E. coli* MG1655 were highly variable (data not shown), whilst in the absence of Pfs, accurate and reproducible levels of MTA were determined (Figure 2c).

Furthermore, higher basal levels of MTA were detected in the *E. coli* Δ*pfs* mutant compared with *E. coli* MG1655. In both genetic backgrounds, there was a clear trend indicating that in the presence of an AHL synthase, MTA accumulated. Induction of *lasI* in the *E. coli* Δ*pfs* mutant resulted in a 26-fold increase in MTA whilst induction of *rhll* caused a 14-fold increase in MTA compared to the equivalent empty vector control.

### Supplementation with exogenously added metabolites partially rescued growth defects imposed by QSSM synthases

As Lasl and Rhll production impedes the growth of *E. coli* and drains away AMC metabolites, the possibility that the exogenous addition of a metabolite that feeds into the AMC could restore growth was investigated. Despite an approximate 14-h delay, exogenous methionine promoted the growth of LasI-producers in a concentration dependent manner (Figure 3), indicating that the fitness cost of QSSM synthases can be partially rescued by replenishing the substrates for these enzymes, and thus the cost is likely to be at least partly energetic. Automated sampling facilitated the measurement of growth throughout the entire growth curve. This necessitated growth in small volumes in a microtitre plate which generated overall kinetics that differed from cultures grown in shaking flasks at larger volumes such as depicted in supplementary Figure 4. Parallel exogenous methionine supplementation in flasks also partially rescued growth defects (data not shown).

**Fig. 3.**
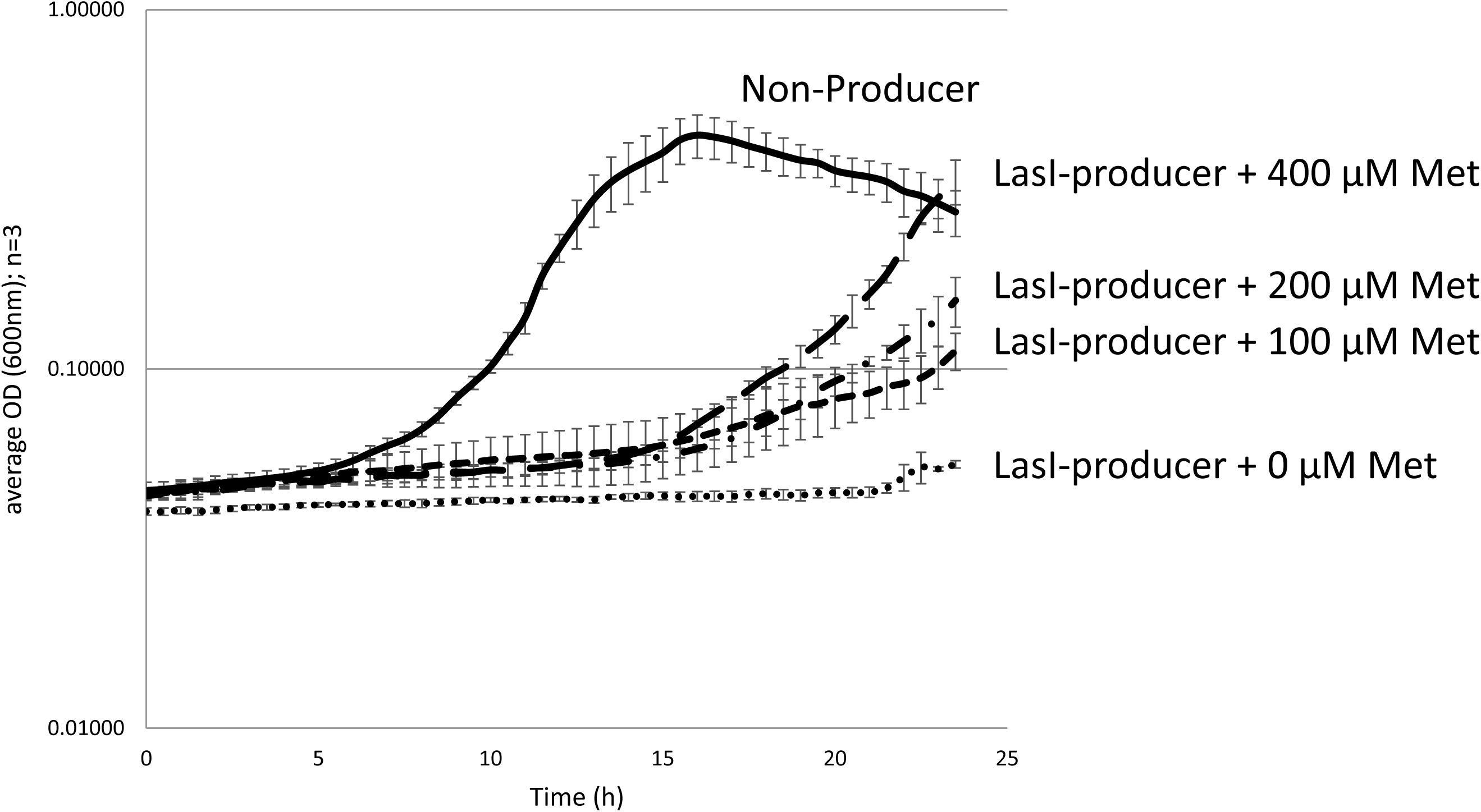
Exogenous methionine addition partially rescues the growth defect imposed by QSSM synthases. MG1655(pME6032) (solid line) and MG1655(pME-*lasl*) (broken lines) were inoculated into MMM + tetracycline, IPTG induced and methionine was added to cultures of MG1655(pME-*lasI*) at time zero. Selected methionine concentrations (μM) are shown. Strains were grown in an automated microplate reader (TECAN Infinite F200) and changes in cell density (OD_600_) monitored. The data are means ± standard deviations for three independent experiments.

### QSSM imposed growth defect confers a fitness cost in mixed populations

Having demonstrated the fitness cost of producing communication signals, we tested whether this was likely to generate a selective advantage in the absence of a beneficial public goods production response, that could cause an evolutionary pressure in conditions more closely mimicking the natural environment where different bacterial strains co-exist in mixed populations. This is particularly important since QSSMs are themselves diffusible public goods available to non-producers. To do this, a QSSM producer and non-producer were grown singly and together, and their relative fitness assessed by ANOVA (Figure 4).

**Fig. 4.**
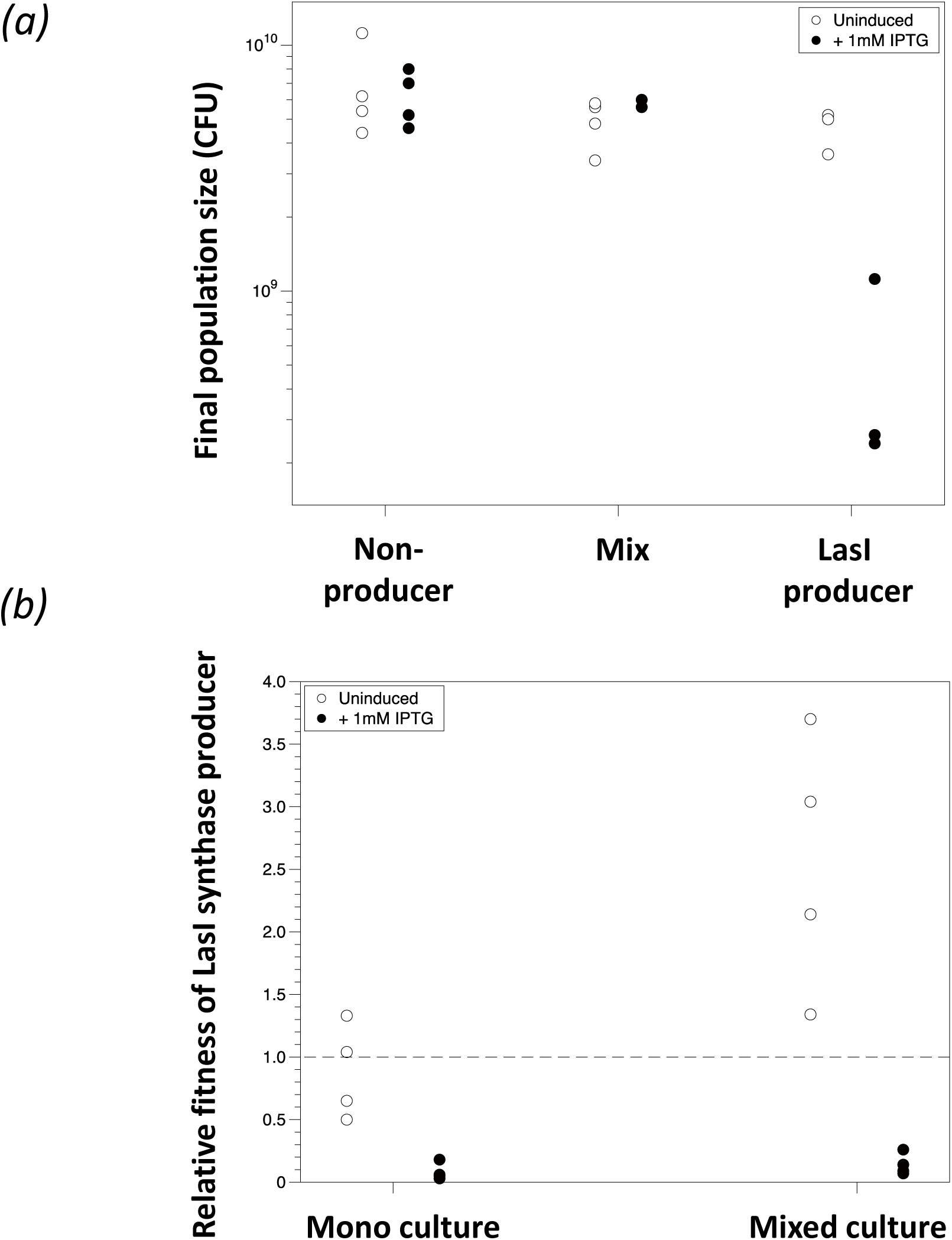
Lasl imposes a fitness cost within a mixed population. *E. coli* MG1655(pME6032), *E. coli* MG1655(pME-*lasl*) or a 1:1 mixture of the two strains were grown. Colony counts were used to calculate the total population density. To determine the population density of each strain in mixed populations, cfu was assessed (**Panel *a***). The evolutionary fitness of *E. coli* MG1655(pME-*lasl*) relative to *E. coli* MG1655(pME6032) in mixed populations is plotted (**Panel *b***). When the two genotypes have equal fitness, *v* = 1. Values of *v* < 1 reflect being outcompeted by MG1655(pME6032 and values > 1 indicate that MG1655(pME-*lasl*) outcompetes MG1655(pME6032). The relative fitness of *E. coli* MG1655(pME-*lasl*) in pure culture was calculated by randomly pairing pure MG1655(pME-*lasl*) and MG1655(pME6032) populations within IPTG treatment.

The total population density was affected by genotype (non-producer, LasI-producer or mix; F_2,18_ = 29.6, *p* < 0.001) and presence/absence of IPTG to induce the AHL synthase (F_1,18_ = 13.4, *p* = 0.002). Moreover, the effect of IPTG depended on population (interaction F_2,18_ = 18.2, *p* < 0.001). As shown in Figure 4a, in the absence of IPTG there were no significant differences in the total densities reached by the non-producer, LasI-producer or mixed populations (Tukey HSD tests, p > 0.47), but in the presence of IPTG the LasI-producer growth was around one log lower than either non-producer or the mix (p < 0.001). Thus adding IPTG decreased growth of LasI-producer, but did not affect the other two populations. Dropping the single outlier from the data set did not affect these results.

The relative fitness of LasI-producer depended on whether the two strains were grown in pure culture or in a mixture (ANOVA: F_1,12_ = 12.9, *p* = 0.004), on the presence of IPTG (F_1,12_ = 78.6, *p* < 0.001) and on the interaction between culture condition and IPTG (F_1,12_ = 6.96, *p* = 0.022). In the absence of IPTG, the two strains grew equally well in pure culture (Figure 4a) and post-hoc *t*-tests showed relative fitness not significantly different from 1: *p* = 0.062) although the LasI-producer had a fitness advantage in mixed culture (Figure 4b) *p* < 0.001). In the presence of IPTG, the relative fitness of LasI-producer was <1 regardless of culture condition (*p* < 0.001).

## Discussion

A number of studies have utilized bacterial QS linked phenotypes to test social evolution theory because QS controls the production of costly ‘public goods’ and as such this creates a drain on the fitness of the cells (Diggle *et al.*, 2007a; Rumbaugh *et al.*, 2009; Kohler *et al.*, 2010; Wilder *et al.*, 2011; Popat *et al.*, 2012; Darch *et al.*, 2012; West *et al.*, 2012; Gupta & Schuster, 2013). Although theory suggests that signals themselves can be costly, there has been no experimental study testing this. Here we provide the first direct evidence that production of the QSSMs themselves, upon which QS relies, is a costly metabolic burden to cells. Specifically we found that (i) bacteria making QSSMs have a growth defect that exerts an evolutionary cost, (ii) production of QSSMs correlates with reduced intracellular concentrations of QSSM precursors, (iii) the production of heteroogous QSSMs negatively impacts the production of a native QSSM that shares common substrates, and (iv) supplementation with exogenously added metabolites partially rescued growth defects imposed by QSSM synthesis‥

Our findings provide experimental support for the theory of Keller and Surrette (Keller & Surette, 2006), who calculated the metabolic cost of QSSM synthesis in terms of ATP. We investigated the cost of signal production at three levels (metabolism, growth, and fitness to compete in co-culture), and showed that the levels of specific central pathway metabolites are altered, and that this has an impact on growth and thus fitness to survive in mixed populations. In the context of horizontal transfer of QS systems, this metabolic perturbation was demonstrated to affect the levels of a native QSSM from the recipient cell suggesting the potential for knock-on effects on the social environment.

A critical question arises regarding the biological relevance of the experimental set up of the study since it uses medium copy (approximately 15) plasmids with an inducible p*trc* promotor rather than a single chromosomal copy of the QSSM synthase encoding genes under the control of their native promoters. The experiments were conducted in this manner to provide us with control over signal production, enabling full induction of the signal at a specified point with the primary aim of determining if signal production can incur a fitness cost. It is possible that even a cost equivalent to a small percentage of what we measured could have a significant impact in natural populations and provide a selective pressure for bacterial evolution. The genes studied here are not naturally plasmid borne, but QS has been extensively studied in this context (Parsek *et al.*, 1997; Cornell & Riscoe, 1998; Parsek *et al.*, 1999; Rumbaugh *et al.*, 1999; Pai *et al.*, 2012; Gupta & Schuster, 2013), as has the activity and impact of many microbial genes including ones that would create a fitness cost that might induce compensatory changes in the native genetic background. Furthermore, the AHLs studied here have been added exogenously to *E. coli* within the context of other studies (e.g. QS reporter strains) without any observable toxic effect on *E. coli*. Moreover, there are plasmid borne QSSM synthase homologues which have been shown to be transferable between bacteria, and presumably this event would impose a fitness burden (Danino *et al.*, 2003; White & Winans, 2007).

Previous work has shown that the major cost to QS is the response to signal (Diggle *et al.*, 2007b). Until now, no one has demonstrated experimentally that the isolated production of the signal itself also incurs a cost. Costly signal production could influence the social environment as it could be subject to social cheating. Self-interest can lead to a breakdown of cooperation at the group level (known as ‘the public goods dilemma’) in bacterial populations, just as it can in animal and human populations (Popat *et al.*, 2012). In environments where QS is important, social cheating on signal production could have important consequences (Brown & Johnstone, 2001). In an infection scenario, it could affect the production of QS-controlled virulence factors. Signal cheating could thus lead to a loss in the ability of bacteria to infect their target hosts (West *et al.*, 2012).

Although *E. coli* contains a protein capable of responding to the production of AHLs (SdiA) (Michael *et al.*, 2001; Yao *et al.*, 2006; Smith *et al.*, 2011; Swearingen *et al.*, 2013), there is no AHL synthase, leading to the notion that *E. coli* can respond to an AHL cue from another species. It is not clear what SdiA regulates in *E. coli*, but it influences cell division, antibiotic resistance and virulence factor production when overproduced. SdiA does not preferentially bind the cognate AHLs for Lasl and Rhll. SdiA has been shown not to interfere with a complete Las QS signalling cassette reporter (Joint *et al.*, 2007; Soares & Ahmer, 2011), and although it can bind to the promoter of *rhll*, it does so regardless of the presence of the cognate AHL (Lindsay & Ahmer, 2005). Importantly, the AHLs detected in our experiments did not overlap significantly with the AHLs shown previously to interact and activate SdiA (Michael *et al.*, 2001; Smith & Ahmer, 2003) and the effect of AHL production upon growth of an *sdiA* mutant mirrored that of MG1655 (data not shown).

Quantification of AHL production, as expected, revealed that the most abundant AHLs were the cognate QSSMs, with pME-*lasI* and pME-*rhll* directing the production of OC_12_-HSL (~30 μM, 71%) and C_4_-HSL (~40 μM, 95%) respectively at the highest IPTG concentration used (supplementary Figure 2c,d). These quantities are broadly in line with AHL production in *P. aeruginosa* where concentrations of 0.5 to 15 μM for OC_12_-HSL and 5-31 μM for C4-HSL have previously been reported (Pearson *et al.*, 1994; Pearson *et al.*, 1995; Cataldi *et al.*, 2008; Ortori *et al.*, 2011).

Such QSSM levels only created a marginal growth defect in rich media supporting our assumption, based on the use of healthy *E. coli* AHL bioreporters, that they would not be toxic (supplementary Figure 4e, 5e). Reduction of signal synthesis, by titration of IPTG, reduced AHLs to undetectable levels and concomitantly repaired the observed growth defects. The extended AHL profile for *E. coli* Rhll-producer was limited to AHLs comprising short acyl chains. In contrast, 9 different long chain AHLs were detected in addition to the cognate OC_12_-HSL in the spent culture supernatants of the LasI-producer, with the quantities of the OC-series dominating. In both cases, the relative lack of leakiness of the IPTG induced promoter was evident by the relatively low levels of AHLs detected in the absence of IPTG. It was not clear why the Rhll inactive mutant S103E did not migrate with the same mobility as all other Rhll proteins analysed, but this may reflect a difference in conformational structure that in turn may influence AHL production by this particular protein.

The observation that growth defects resulting from the introduction of AHL production were more prominent in MMM supports our hypothesis that the metabolic drain of AHL production would occur via the AMC since previous studies have shown that disruption of the AMC is more readily reflected in growth defects in defined media limited for the sulphur sources that feed into this pathway (Winzer *et al.*, 2003; Doherty *et al.*, 2006; Heurlier *et al.*, 2009; Holmes *et al.*, 2009; Doherty *et al.*, 2010).

The approximate energy cost of AHL production was crudely calculated. It has been estimated that an *E. coli* cell contains 12.1 billion ATP molecules (Stouthamer, 1973). Using a value of 30 μM corresponding to the highest concentration of OC_12_-HSL observed at 1 mM IPTG, we estimate the total cost of ATP production to be a significant energetic cost at ~7% of the ATP.

This is in line with metabolic perturbations of QS (Davenport, P., Griffin, J.L., Welch, M., 2015), however has to be further investigated experimentally to accurately reflect the metabolite turnover in the conditions studied. The ability of the Rhll-producer to achieve the same final population density as the non-producer may reflect a delay in the collective benefit of QS. The production of QS-regulated public goods does not instantly exceed the cost of QSSM production. Once the population density reaches a threshold, the associated high level of QSSMs may provide a greater benefit in the context of QS regulated public goods production (Pai *et al.*, 2012). Interestingly, measurement of population density by OD in Figure 1 indicated an approximate 10-fold fitness advantage in monoculture compared to the approximate 4-fold change in relative fitness calculated for Figure 4 between non-producer and LasI-producer using viable cells (CFU). It was notable that in mixed culture (Figure 4), the faster growth of the non-producer did not exhaust the overall nutrient supply and thereby greatly reduce the fitness estimate of LasI-producer in mixed culture compared to that in monoculture. It is not clear what the underlying reason for this is, or why Lasl-producer is fitter than the control in the uninduced mixed culture compared to the uninduced monoculture in Figure 4 given that *E. coli* is incapable of responding to AHL signals and thus not participating in a compensatory production of beneficial public goods. This observation suggests that LasI-producer benefits from the presence of the other strain. There is a wide spread of relative fitness estimates for the uninduced mixed culture over the 4 parallel samples which may be reduced through inclusion of further replicas, but it is also possible that the control strain reaches stationary phase quickly, and some of the population lyses releasing nutrients that the LasI-producer can use to grow on, leading to a rise in viable cells. Whatever the underlying reason, this finding accentuates the main finding of this experiment: that producing the AHL synthase is costly when in the context of a mixed culture. It would be interesting to extend this approach by determining the relative fitness of the different AHL-producing mutants that we generated and also to determine whether Rhll-producers (that begin growing slowly but ultimately reach densities equal to non-producers in monoculture) are disadvantaged in mixed populations. Extending these studies would also facilitate measuring rates of ATP/signal turnover per cell during growth to enable calculations of the energetic cost of signal production to relate to theoretical values.

Interestingly, despite the LasI protein being produced at a higher level, the total level of AHLs and the level of the cognate AHL synthesized by LasI were similar to those made by Rhll (supplementary Figure 2, 4a and 5a). The greater effect this had upon growth is likely to result in part from the need for the longer acyl chain on the AHLs made by LasI (12 carbons compared to 4 carbons on the cognate AHLs of Lasl and Rhll respectively). This would require greater metabolic investment from fatty acid biosynthesis pathways and thereby invoke a greater fitness cost. Despite the similar overall AHL levels, which would be predicted to require similar levels of the other shared substrate (SAM) and thus other AMC-related metabolites, there was a more substantial drain on SAM and other AMC metabolites that led to a corresponding fall in AI-2 and rise in MTA levels which in turn could influence polyamine levels since MTA is a feed-back inhibitor of polyamine synthesis. The reason for this requires further investigation, but could reflect changes in metabolic flux through the AMC triggered by the relative demands on fatty acid metabolism which are linked via cysteine (which feeds into coenzyme A biosynthesis and also the AMC).

The correspondence between the higher level of LasI production and a more dramatic effect on growth and metabolite levels, offers the potential to titrate in effects on fitness gradually by manipulating levels of AHL synthase production. This enhances the potential of using QS bacteria as a model to test aspects of signalling theory by experimentally manipulating the cost (Smith & Harper, 2003). Furthermore, revealing that the production of AHLs perturbs the levels of central metabolites within bacteria and has a knock-on effect on cell fitness provides impetus to drive the development of novel medical intervention strategies given the contribution of QS to virulence. The design of future therapeutic agents may include the previously proposed exploitation of QS by free-loaders (bacteria that neither produce nor respond to QSSMs) to reduce population size and virulence (Allen *et al.*, 2014). Another alternative would be to exploit the finding here that the fitness cost of QSSM production is dependent upon the supply of nutrients. As proposed in (Hall *et al.*, 2011), administering QSSMs alongside the antibiotic rifampicin would induce the expression of many genes including the QSSM synthases, thereby creating an increased demand for RNA polymerase and elevating the cost of rifampicin resistance. Such constraints on the evolution of resistance may prolong the utility of antibiotics, both current and future.

## Acknowledgements

We thank Alex Truman for the synthesis and provision of QSSMs. This study was supported by funds from the European Union, Biotechnology and Biological Sciences Research Council, Natural Environment Research Council, Libyan Government and Royal Society.

## Supplementary Material

**Supplementary Figure Legends**

**Supplementary Fig. 1.**
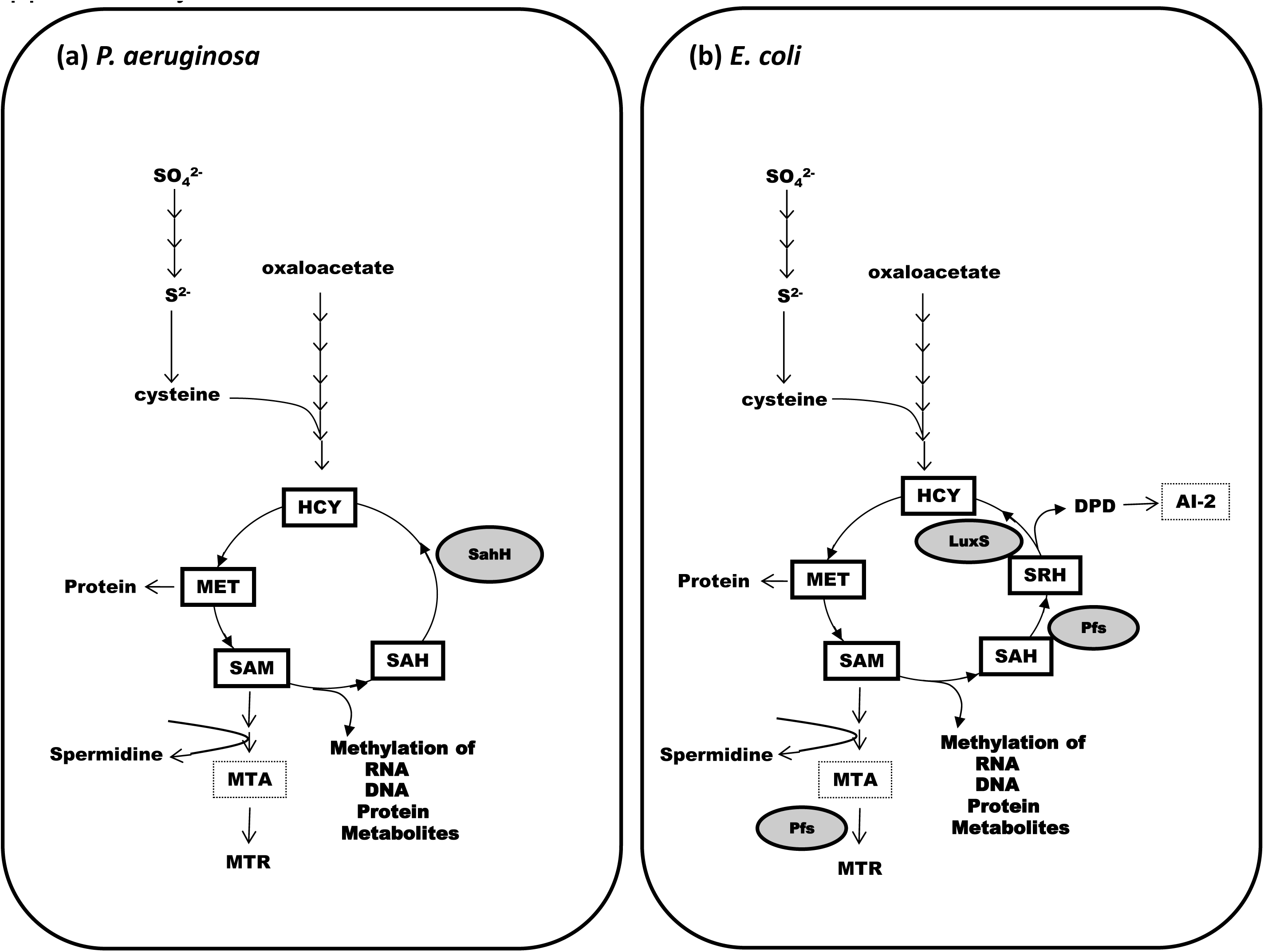
The activated methyl cycle of (a) *P. aeruginosa* and (b) *E. coli*. In *P. aeruginosa* SAH hydrolase (SahH) converts SAH (*S*-adenosyl homoserine) to HCY (homocysteine) directly, but in *E. coli* the enzymes Pfs and LuxS work sequentially generating the intermediate SRH (*S*-ribosyl homoserine) and the additional product DPD (4,5-dihydroxy-2,3-pentanedione) which is converted to AI-2 (autoinducer 2). In *E. coli*, Pfs has a secondary role in conversion of MTA (5’-methylthioadenosine) to MTR (5’-methylthioribose). MET: methionine, SAM: *S*-adenosyl-L-methionine.

**Supplementary Fig. 2.**
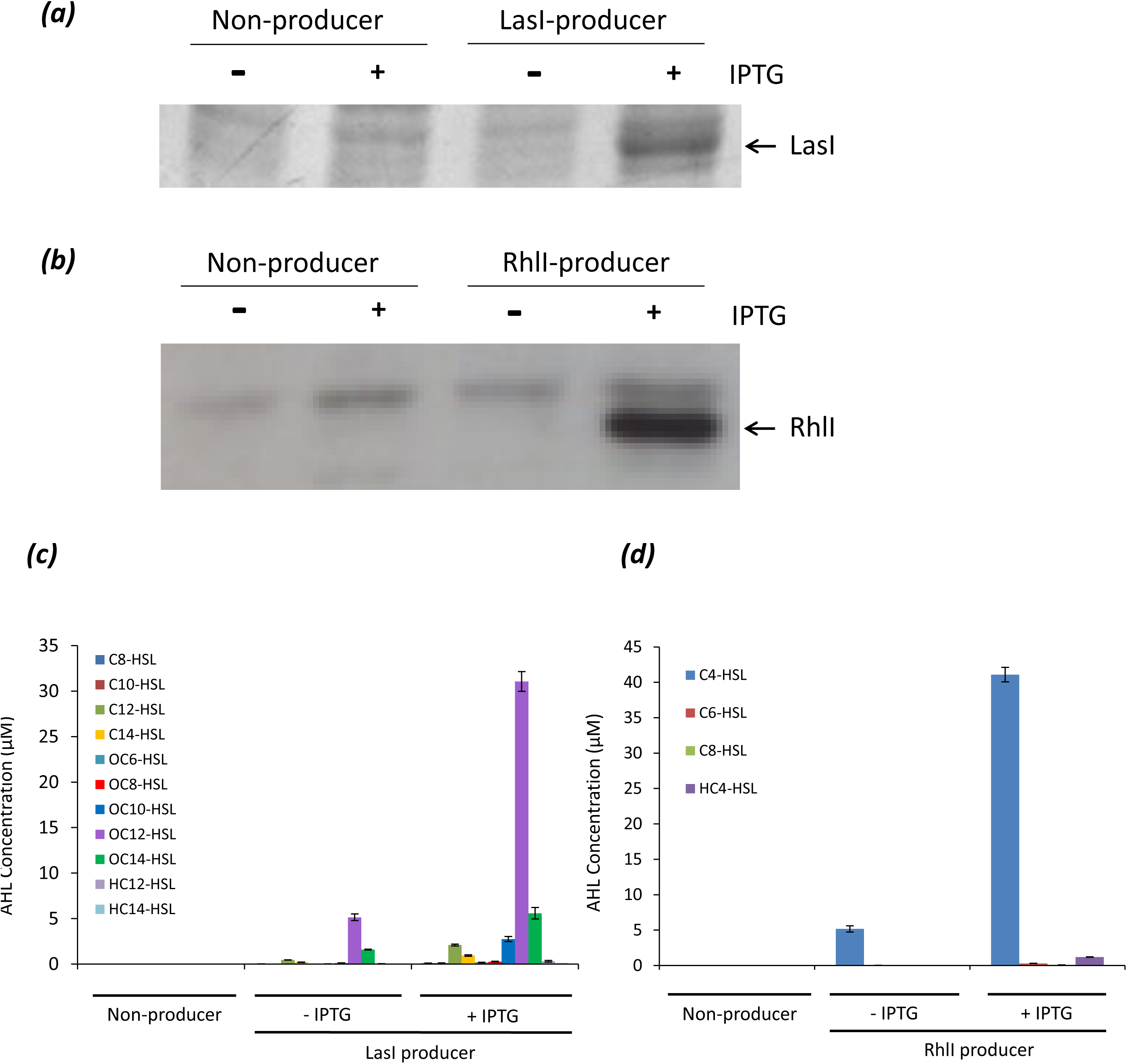
AHL synthases direct the synthesis of their cognate QSSMs in a heterologous host. *E. coli* strains MG1655[pME6032], MG1655[pME-*rhll*] and MG1655[pME-*lasI*] were grown in LB + tetracycline until an OD_600_ = 0.5 and induced with 1 mM IPTG for 2 h. Whole cell extracts were separated by SDS PAGE and the presence of LasI **(panel *a*)** was detected by Coomassie staining whist the lower levels of RhlI were detected by Immunoblotting with a specific antisera **(panel *b*)**. Quantitative profiling of AHLs produced by the same strains was undertaken by extracting with acidified ethyl acetate from LB or late exponential phase supernatants of *E. coli* strains MG1655[pME6032] (OD_600_ = 0.8), MG1655[pME-*lasI*] (OD_600_ = 0.9), and MG1655[pME-*rhll*] (OD_600_ = 0.9) grown in LB containing IPTG. The actual concentration (μM) determined by LC-MS/MS analysis of AHLs described in Experimental Procedures for *lasI* **(c)** and *rhll* **(*d*)** is shown. Of the 11 AHLs detected in cultures harvested from *lasI* induced cells OC_10_-HSL, OC_12_-HSL and OC_14_-HSL were the dominant signals. Other AHLs present below 5 μM were C_8_-HSL, C_10_-HSL and C_12_-HSL, C_14_-HSL, OC_6_-HSL, OC_8_-HSL, HC_12_-HSL and HC_14_-HSL. The 4 AHLs detected in culture supernatant of MG1655[pME-*rhll*] were C_4_-HSL, C_6_-HSL, C_8_-HSL and HC_4_-HSL. The data are means ± standard deviations for three independent extractions.

**Supplementary Fig. 3.**
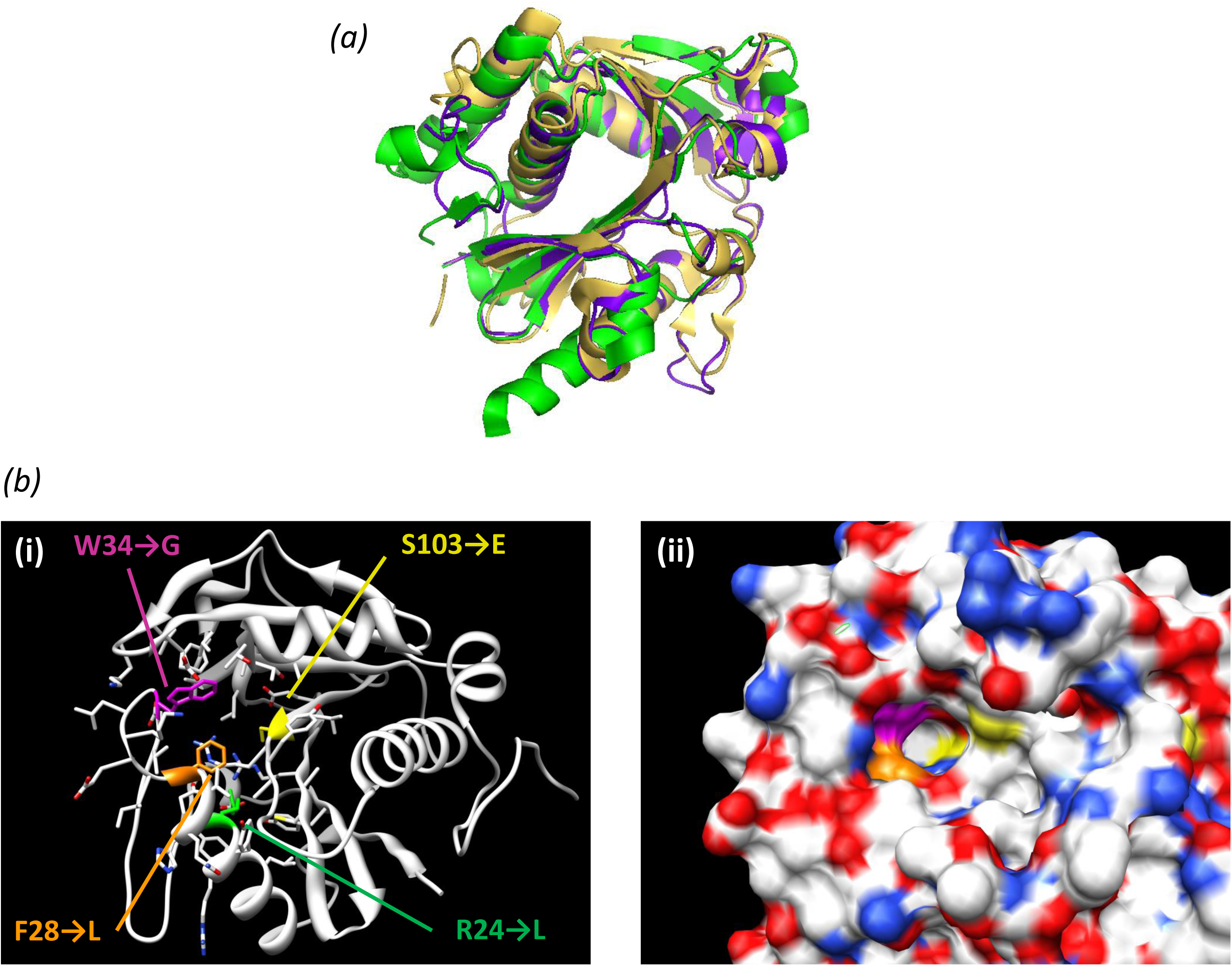
Rhll modelling to inform the creation of catalytically inactive QSSM synthase mutants. *P. aeruginosa* Rhll model (purple) was predicted using multi-template homology modelling, and utilised the structures of *P. aeruginosa* LasI (yellow) and *P. stewartii* EsaI (green) **(*a*)**. The positions of the Lasl and Rhll conserved residues selected for mutation in this study are shown on the ribbon **(*b*)** and space filling **(*c*)** models on the predicted structure of Rhll. Colours indicate R23 (green), F28 (orange), W34 (pink) and S103W (yellow). **Rhll Homology Modelling**. An atomised model of Rhll was constructed using MODELLER9v7 [31] and the homologs *P. aeruginosa* LasI (30% identity with Rhll sequence; pdb code 1r05) and *P. stewartii* EsaI (22% identity with Rhll sequence; pdb code 1kzf) as templates. The model with the lowest DOPE score [32] was chosen as the best model and used for visualization of the Rhll protein. The model was checked and hydrogen atoms added/refined using MOLPROBITY [33]. The resulting ramachandran plot demonstrated that 92.5% of model residues were in favoured regions.

**Supplementary Fig. 4.**
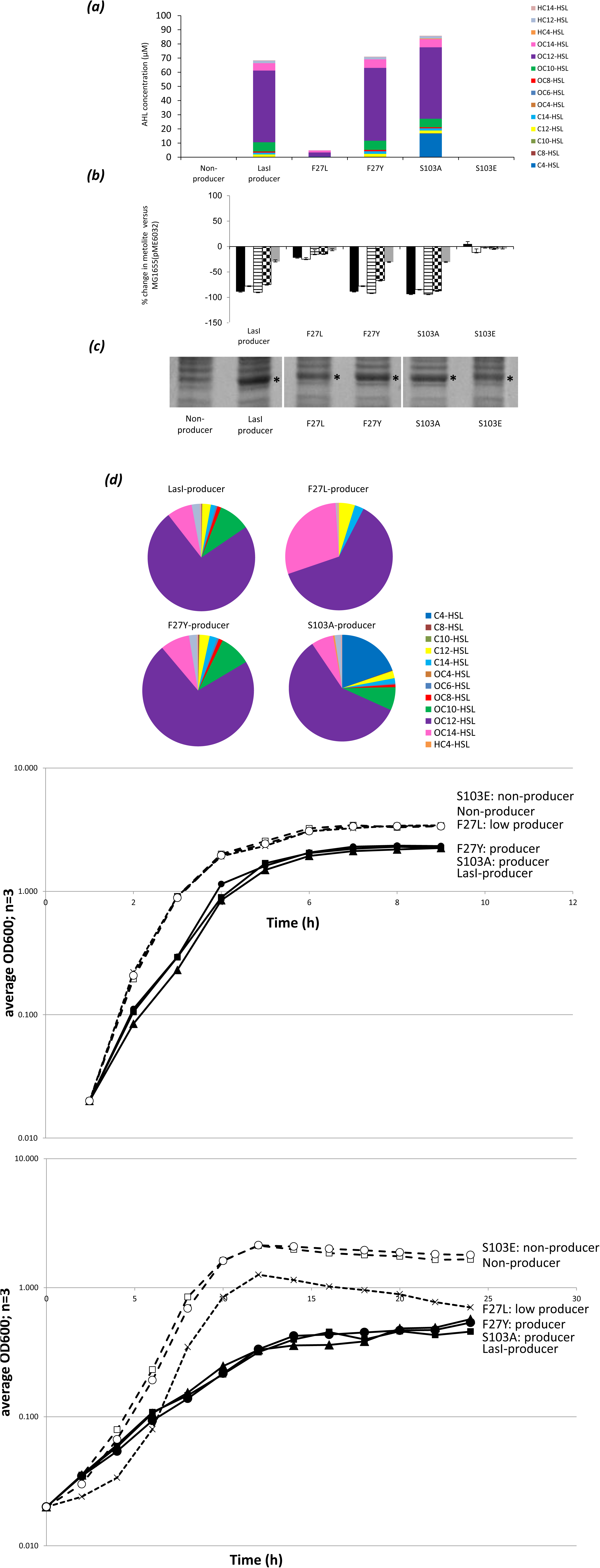
QSSM synthase mutants lacking the ability to generate AHLs no longer impose a fitness cost and fail to make an impact on the levels of AMC metabolites: LasI. *E. coli* MG1655 bearing the empty vector (pME6032) or a derivative encoding WT LasI (pME-*lasl*), or LasI mutated to introduce the change F27L, F27Y, S103A, or S103E, were grown in the presence or absence of IPTG as indicated. The production of LasI was monitored by Coomassie staining the SDS PAGE (**panel (*c*)**, marked with an asterix). Size was estimated by comparison with the molecular weight markers. **Panel (*a*)** shows the AHLs extracted from late exponential phase supernatants of each strain with acidified ethyl acetate and profiled by LC-MS/MS analysis as described in Experimental Procedures. The ratio between the different signalling molecules present in supernatants is represented by pie charts in **Panel (*d*)**. The number of repeats represented is 3. **Panel (*b*)** Metabolite levels were determined in the same strains following growth in LB with IPTG induction until OD 0.8-0.9. Intracellular accumulation of SAM (black), SAH (white), SRH (horizontal stripes), HCY (checked) and MET (grey) was determined by LC-MS analysis. The peak area corresponding to each compound in an extract was divided by the peak area of an appropriate internal standard (IS) for normalisation; the data are the means ± standard deviations for three independent cultures. The same *E. coli* SDM strains were inoculated into 125 ml LB + tetracycline media **Panel (*e*)** or MMM + tetracycline **Panel (*ƒ*)** and grown shaking at 37°C in 500 ml-Erlenmeyer flasks. Aliquots (1 ml) were taken at regular intervals as indicated, and the mean OD_600_ values of triplicate culture samples are shown on a log_10_ scale over time (h). Error bars indicate standard deviations from the means. Strains generating wild type levels of AHLs are indicated by the solid lines and closed symbols (pME-*lasl*, *lasl*S103A, *lasl*F27Y), those not making detectable AHLs by the dashed lines and open symbols (pME6032, *lasl*F27L, *lasl*S103E), and those producing intermediate levels by the dotted lines with cross symbols (MG1655(pME-*lasl*F27L)). The MG16 (pME6032) negative control is indicated by open squares and the positive control by closed squares: MG1655(pME-*las*I). The symbols used in the figure are closed triangles for the *lasl*F27Y, closed circles for *lasl*S103A, open circles for *lasl*S103E.

**Supplementary Fig. 5.**
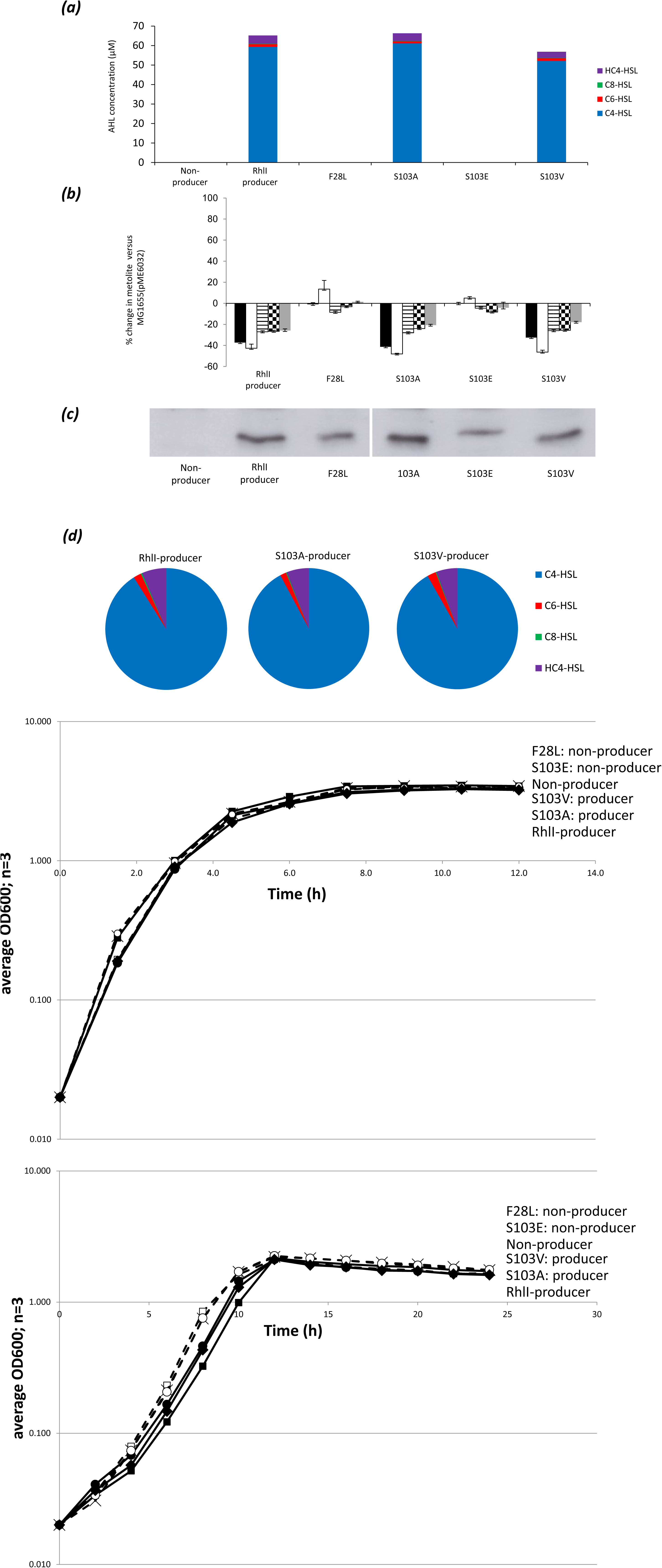
QSSM synthase mutants lacking the ability to generate AHLs no longer impose a fitness cost and fail to make an impact on the levels of AMC metabolites: Rhll. *E. coli* MG1655 bearing the empty vector (pME6032) or a derivative encoding WT Rhll (pME-*rhll*), or Rhll mutated to introduce the change F28L, S103A, S103E, or S103V were grown in the presence or absence of IPTG as indicated. The production of Rhll was monitored by immunoblotting with anti-Rhll (**panel (*c*)**). Size was estimated by comparison to the molecular weight markers. **Panel (*a*)** shows the AHLs extracted from late exponential phase supernatants of each strain with acidified ethyl acetate and profiled by LC-MS/MS analysis as described in Experimental Procedures. The ratio between the different signalling molecules present in supernatants is represented by pie charts in **Panel (*d*)**. This was repeated 3 times. **Panel (*b*)** Metabolite levels were determined in the same strains following growth in LB + tetracycline with IPTG induction until OD 0.8-0.9. Intracellular accumulation of SAM (black), SAH (white), SRH (horizontal stripes), HCY (checked) and MET (grey) was determined by LC-MS analysis. The peak area corresponding to each compound in an extract was divided by the peak area of an appropriate internal standard (IS) for normalisation; the data are the means ± standard deviations for three independent cultures. The same *E. coli* SDM strains were inoculated into 125 ml LB + tetracycline media **Panel (*e*)** or MMM + tetracycline **Panel (*ƒ*)** and grown shaking at 37°C in 500 ml-Erlenmeyer flasks. Aliquots (1 ml) were taken at regular intervals as indicated, and the mean OD_600_ values of triplicate culture samples are shown on a log_10_ scale over time (h). Error bars indicate standard deviations from the means. Strains generating wild type levels of AHLs are indicated by the solid lines and closed symbols (pME-*rhll*, *rhll*S103A, *rhll*S103V), those not making detectable AHLs by the dashed lines and open symbols (pME6032, *rhll*F27L, *rhll*S103E). The MG1655(pME6032) negative control is indicated by open squares and the positive control by closed squares: MG1655(pME-*rhll*). The symbols used in the figure are closed diamonds for the *rhll*S1-3V, closed circles for *rhll*S103A, open circles for *rhll*S103E.

**Supplementary Fig. 6.**
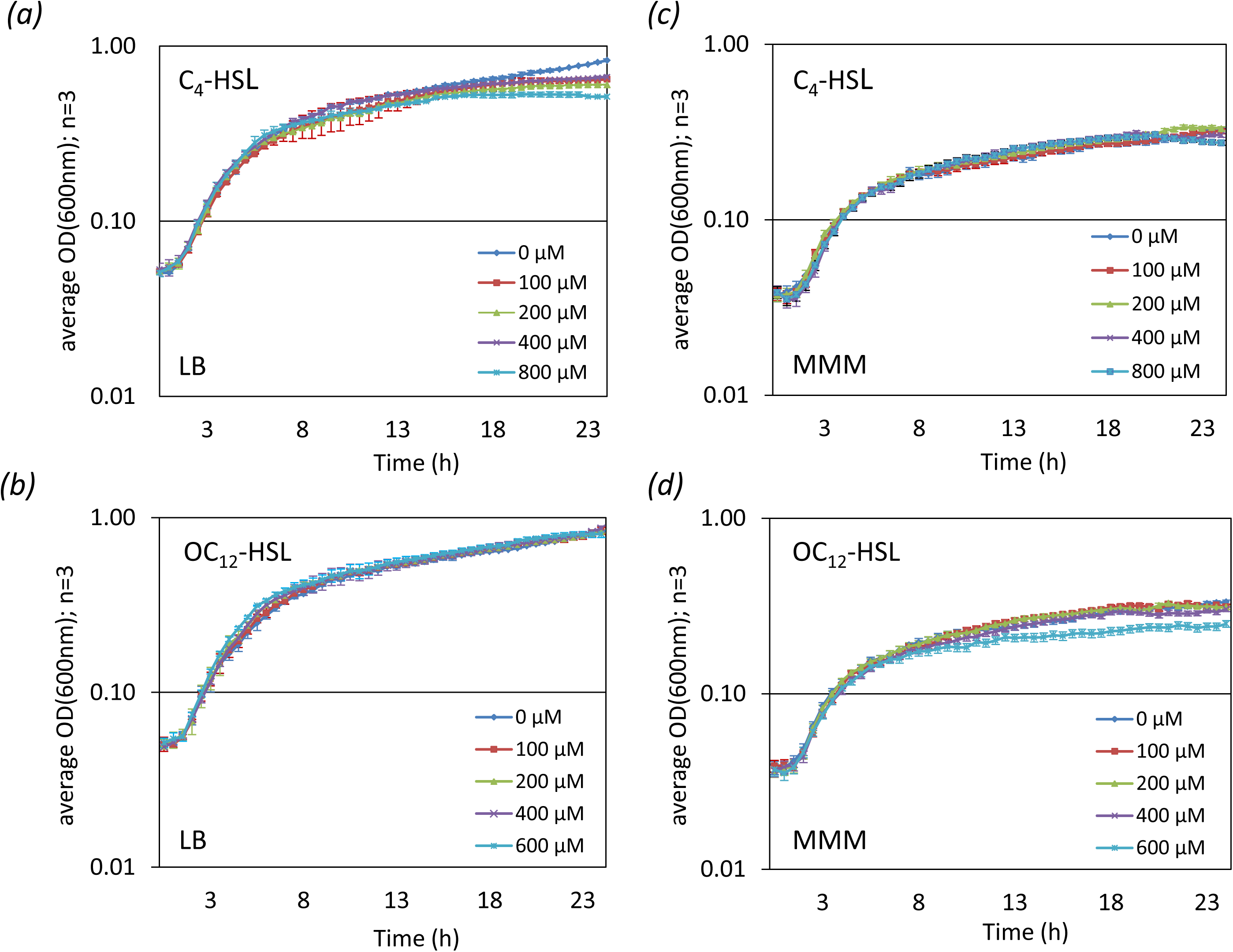
Exogenous addition of QSSMs does not affect growth of a heterologous host. *E. coli* strain MG1655[pME6032] was grown with varying concentrations of C4-HSL or OC12-HSL added exogenously into LB (a, b) or MMM cultures (c, d) at the start of the experiment. The optical density was determined during growth every 30 min at wavelength of 600 nm using a TECAN microplate reader. Standard deviations are based on the mean values of three parallel cultures.

---

**Supplementary Table 1.**
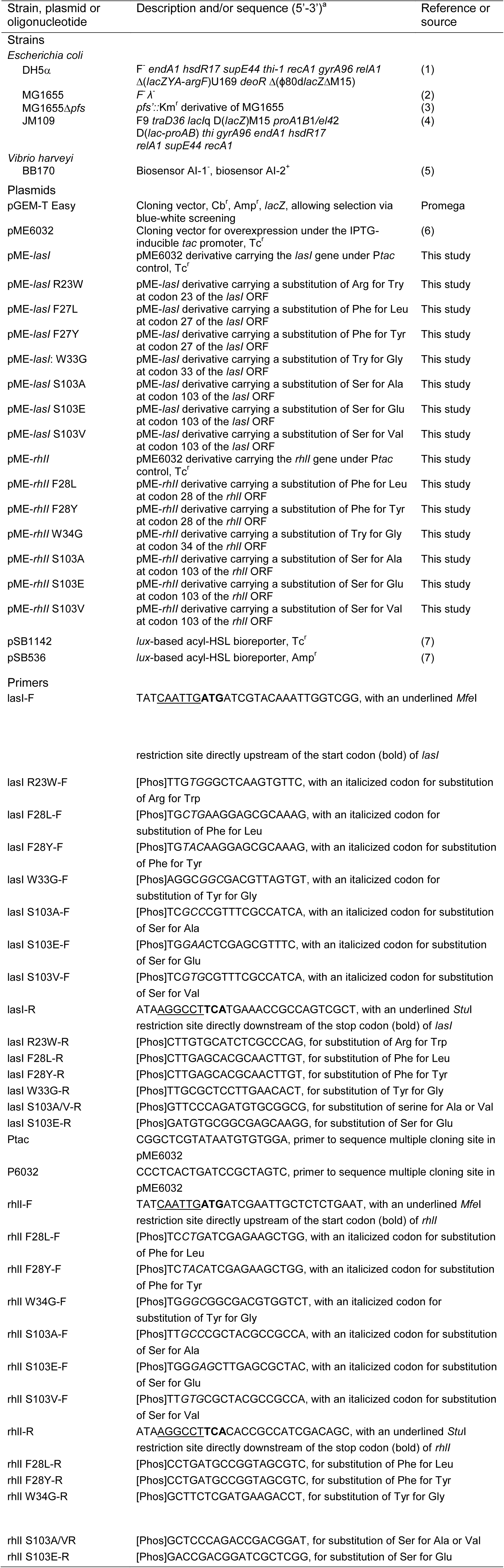
**Strains, plasmids and primers used in this study**.

**Supplementary Table 2.** Substitutions chosen for mutagenesis of AHL synthases. Residues chosen aligned with those published in (1), and map to equivalent positions in *lasI*. Other substitutions chosen were predicted to insert more structurally conservative changes.

| AA position | AA Encoded | WT DNA sequence | Mutant AA | Mutant DNA sequence |
| --- | --- | --- | --- | --- |
| *lasI* |  |  |  |  |
| 23 | R | CGT | W | TGG |
| 27 | F | TTC | L | CTG |
|  |  |  | Y | TAC |
| 33 | W | TGG | G | GGC |
| 103 | S | AGC | A | GCC |
|  |  |  | E | GAA |
|  |  |  | V | TGC |
| *rhII* |  |  |  |  |
| 28 | F | TTC | L | CTG |
|  |  |  | Y | TAC |
| 34 | W | TGG | G | GGC |
| 103 | S | TTC | A | GCC |
|  |  |  | E | GAG |
|  |  |  | V | GTG |

**Supplementary Table 3.**
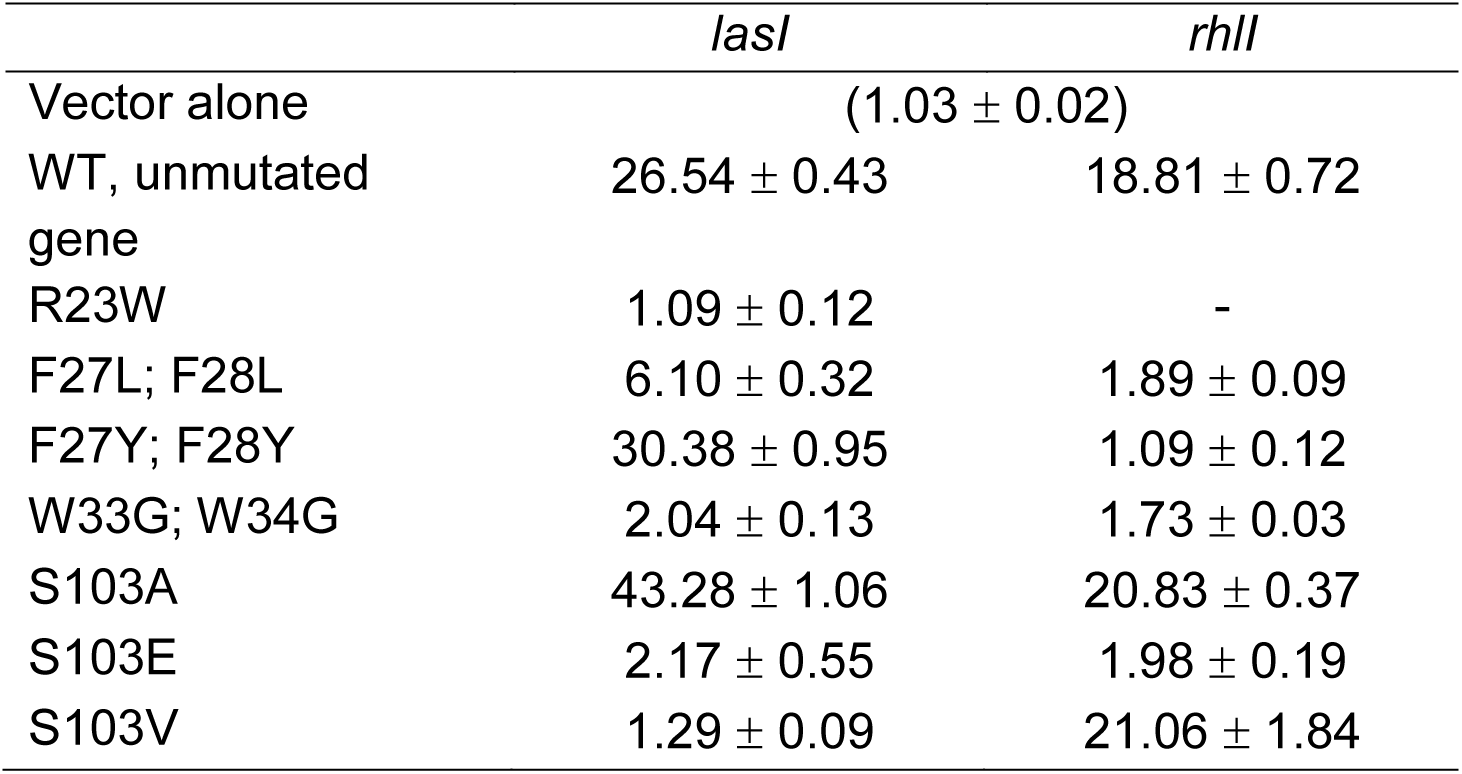
MTA levels in strains bearing QSSM synthases. Figures for metabolites attained in one representative independent experiment.

^†^Concentrations (in relative units) of AMC metabolites were determined by analysing cell content using LC-MS analysis, as described in Materials and Methods. The mean OD_600_ values of triplicate culture samples standard deviations from the means are shown.

**Supplementary Table 4.** Intracellular concentrations of AMC metabolites decrease in *E. coli* producing active QSSM synthases. Figures for metabolites attained in one representative independent experiment.

| Met. | Cell density (OD<sub>600</sub>) | MG1655 [pME6032] –IPTG | MG1655 [pME6032] +IPTG | MG1655 [pME-*lasI*] –IPTG | MG1655 [pME-*lasI*] +IPTG | MG1655 [pME-*rhlI*] –IPTG | MG1655 [pME-*rhlI*] +IPTG |
| --- | --- | --- | --- | --- | --- | --- | --- |
| SAM |  | 39.46 ± 0.72 | 45.67 ± 1.11 | 43.85 ± 2.25 | 1.39 ± 0.02 | 39.77 ± 0.86 | 23.40 ± 0.29 |
| SAH |  | 0.109 ± 0.001 | 0.112 ± 0.010 | 0.102 ± 0.004 | 0.029 ± 0.004 | 0.086 ± 0.002 | 0.058 ± 0.006 |
| SRH |  | 0.590 ± 0.025 | 0.561 ± 0.016 | 0.564 ± 0.007 | 0.060 ± 0.004 | 0.599 ± 0.051 | 0.444 ± 0.064 |
| HCY |  | 8.204 ± 0.819 | 8.250 ± 0.486 | 8.614 ± 0.291 | 2.109 ± 0.268 | 8.834 ± 1.360 | 6.360 ± 0.216 |
| MET |  | 188.539 ± 5.668 | 182.668 ± 11.185 | 180.537 ± 3.735 | 127.390 ± 4.525 | 185.038 ± 12.581 | 146.284 ± 4.002 |

^†^Concentrations (in relative units) of AMC metabolites were determined by analysing cell content using LC-MS analysis, as described in Materials and Methods. The mean OD_600_ values of triplicate culture samples standard deviations from the means are shown.

